# Do anglers take the bait? Anglers’ perceptions about fluvial barriers in three river basins in Northern Spain

**DOI:** 10.64898/2026.02.18.706343

**Authors:** Ana Sánchez-Alcázar, Rafael Miranda, David Galicia, Lide de Izeta-Zalduendo, José Barquín, Alexia M. González-Ferreras, Francisco J. Peñas, Ana Villarroya

**Affiliations:** Universidad de Navarra, Institute for Biodiversity and Environment (BIOMA), Irunlarrea 1, 31008 Pamplona, Spain; Universidad de Navarra, Faculty of Sciences, Department of Environmental Biology, Environmental Humanities (HUMAM), Irunlarrea 1, 31008 Pamplona, Spain; Universidad de Navarra, Faculty of Sciences, Department of Environmental Biology, Biodiversity Data Analytics and Environmental Quality (BEQ), Irunlarrea 1, 31008 Pamplona, Spain; IHCantabria – Institute of Environmental Hydraulics, University of Cantabria, PCTCAN, C/Isabel Torres 15, 39011 Santander, Cantabria, Spain

**Keywords:** angling, dam removal, river restoration, social perception, Spain, telephone survey, weir removal

## Abstract

Removing river barriers - such as dams or weirs - is an increasingly used strategy for restoring freshwater ecosystems. In Europe, these actions are key for achieving the goal of 25,000 kilometers of free-flowing rivers that the recent regulation on nature restoration establishes for 2030. However, social acceptance remains uneven, and local opposition—often related to cultural attachments, poor ecological awareness, and misinformation—may influence or even impede restoration efforts. Among stakeholders, anglers play a particularly influential role, yet their perceptions and knowledge remain poorly documented. This study addresses how anglers from three river basins in Northern Spain perceive river barriers, their removal, and their ecological impacts, and contrasts their attitudes to those of other residents. We carried out a telephone survey of 1,200 adult residents in the target basins. We assessed perceptions, misconceptions, and self-reported knowledge of river barriers, and collected various sociodemographic parameters. We selected 180 self-identified anglers and compared their answers to those from a subsample of 180 non-anglers with similar demographic characteristics. Despite reporting significantly higher self-perceived knowledge and more polarised responses, anglers showed lower awareness of the ecological impacts of fluvial barriers. They were more likely to underestimate their negative effects compared to the control group. In addition to falling for the main misconceptions surrounding the issue, their responses displayed a degree of bimodality, suggesting that the type of fishing practised may influence their attitudes. Our findings reaffirm the importance of strengthening awareness-raising efforts among relevant stakeholders about the impacts of river barriers and the benefits of their removal when planning specific interventions. It is essential to engage local communities—particularly key groups such as anglers—to strengthen the social acceptance of such actions and improve environmental governance.

## Introduction

Rivers, essential components of the planet’s ecological network and key elements in the development of human societies, are currently subject to numerous impacts and pressures, among which the presence of multiple artificial transversal barriers along their courses stands out (Grill et al., 2019; Belleti et al., 2020).

Dams, weirs, and other barriers have become highly useful infrastructures for managing water resources, enabling the storage and direct use of water for domestic supply, agriculture and industrial uses, as well as for flood control and hydroelectric power generation (Richter et al., 2010; Perera & North, 2021). Beyond their primary function, even when these structures are no longer in use, they often provide recreational opportunities associated with activities such as hiking, fishing, boating or swimming and can also create visually and acoustically appealing waterscapes that may become part of the cultural heritage and identity of the surrounding area (Born et al., 1998; Brummer et al., 2017). Nevertheless, the fragmentation these constructions cause in river systems has numerous negative ecological consequences (Barbarossa et al., 2020; He et al., 2024; Chan et al., 2025).

Upstream of these barriers, the river acquires the characteristic dynamics of lentic aquatic ecosystems due to a reduction in water velocity because of water storage and the flooding of adjacent areas. This process can alter riparian forests, the groundwater table, and water physicochemical properties, including temperature, dissolved oxygen, pH, and conductivity, while also leading to the accumulation of sediments and pollutants (Nilsson et al., 2005; Zang et al., 2024). Downstream, the fluvial dynamics are modified by the alteration of the natural flow, sediments, temperature, nutrient and materials, which may have detrimental effects not only on river ecosystems, including river channels, riparian zones and floodplains, but also on estuarine and even coastal areas (Nilsson & Berggren, 2000; Richter et al., 2010, Peñas et al., 2014).

These alterations have numerous consequences, particularly for aquatic flora and fauna, as longitudinal connectivity, habitat conditions and resources availability are significantly modified. In Europe, migratory fish species as emblematic as Atlantic salmon (*Salmo salar*), brown trout (*Salmo trutta*), or European eel (*Anguilla anguilla*), are especially affected, losing access to breeding and spawning riverbeds essential for completing their life cycles (Liermann et al., 2012; González-Ferreras et al., 2019; Waldman & Quinn, 2022). Many other aquatic species also experience population fragmentation and reduced survival rates (Van Looy et al. 2014; Seliger & Zeiringer, 2018), including plant species that alter their distribution, community composition and seed dispersal patterns (Bunn & Arthington 2002). Furthermore, eutrophication phenomena (Ma et al., 2023) and the proliferation of invasive species become more frequent in river systems affected by these obstacles (Fritts et al., 2024).

In view of these consequences, the European Green Deal (EC, 2019), the EU Biodiversity Strategy for 2030 (EC, 2020) and Nature Restoration Law (Regulation (EU) 2024/1991) advocate, among their measures, the removal of obsolete barriers from river channels to restore fluvial ecosystems and enhance their resilience to future climate scenarios. However, when such restoration measures are undertaken, local opposition frequently arises, as these barriers have often become part of local identity or continue to provide recreational benefits to communities nearby (Lejon et al., 2009). In addition, there is a widespread lack of environmental awareness regarding their adverse effects and an increasing politicisation and misinformation surrounding the issue (Fox et al., 2016; Reilly & Adamowski, 2017; Hommes, 2022). As a consequence, planned river restoration projects can generate social discontent and unpopularity, and, in some cases, fail to be implemented altogether (Diessner et al., 2020; Parent et al., 2024).

Knowing river users’ perceptions is essential to better understand phenomena of opposition or lack of awareness, and to provide local administrations with reliable information to address obstacles to removal processes and to support awareness campaigns (Johnson & Graber, 2002). Even if international studies provide valuable data in this regard, social perception is highly influenced by cultural contexts. Currently, there is a lack of knowledge about the social perception of dams and weirs in Europe (Dopico et al., 2022), a knowledge gap that must be addressed if we want to carry out EU fluvial restoration activities efficiently and soundly.

In this context, assessing the role that anglers, perceptions might have on river connectivity restoration projects is paramount, as they constitute one of the largest and most influential cultural groups directly linked to river ecosystems (van den Heuvel & Rönnbäck, 2023; Banha et al., 2024), whose involvement in dam removal processes is recurrent (Lejon et al., 2009; Boucher and Hudson, 2023). For instance, in Spain, anglers currently hold more than 500,000 fishing permits, with 30,000 of these licences issued in the autonomous communities within the study area (Ministerio para la Transición Ecológica y el Reto Demográfico, 2025). In fact, Fishers’ Ecological Knowledge (FEK) is increasingly being used as a complementary source of information for environmental management (Eden, 2012; Bevilacqua et al., 2016), although not much research has been done to assess their specific perspective or level of knowledge on the effects of built barriers in rivers.

To address the gap, the main goal of this study is to evaluate anglers’ perceptions and awareness of river barriers and their removal, with the aim of informing the efficient management of removal processes and river conservation. Specifically, this study examines three main research questions: 1) How do anglers perceive these barriers and their removal? 2) Are anglers aware of the barriers’ ecological impacts? 3) Do anglers’ perceptions of these issues differ from those of other locals who do not fish? In light of these questions, we hypothesise that anglers’ perceptions of fluvial obstacles, their removal, and the associated ecological impacts will differ from those of the general public in the study region, primarily in that anglers are more aware and precise.

## Methods

### Study Area

The study addresses the case of northern Spain, a territory with a high density of dams per kilometre of river in Europe (Belletti et al., 2020) that also holds one of the highest percentages of endemic freshwater fish species in that region (Reyjol et al., 2007; Miqueleiz et al., 2022). Within that area, the research focuses on three Atlantic basins: Deva-Cares, Oria, and Bidasoa (Figure 1).

**Figure 1.**
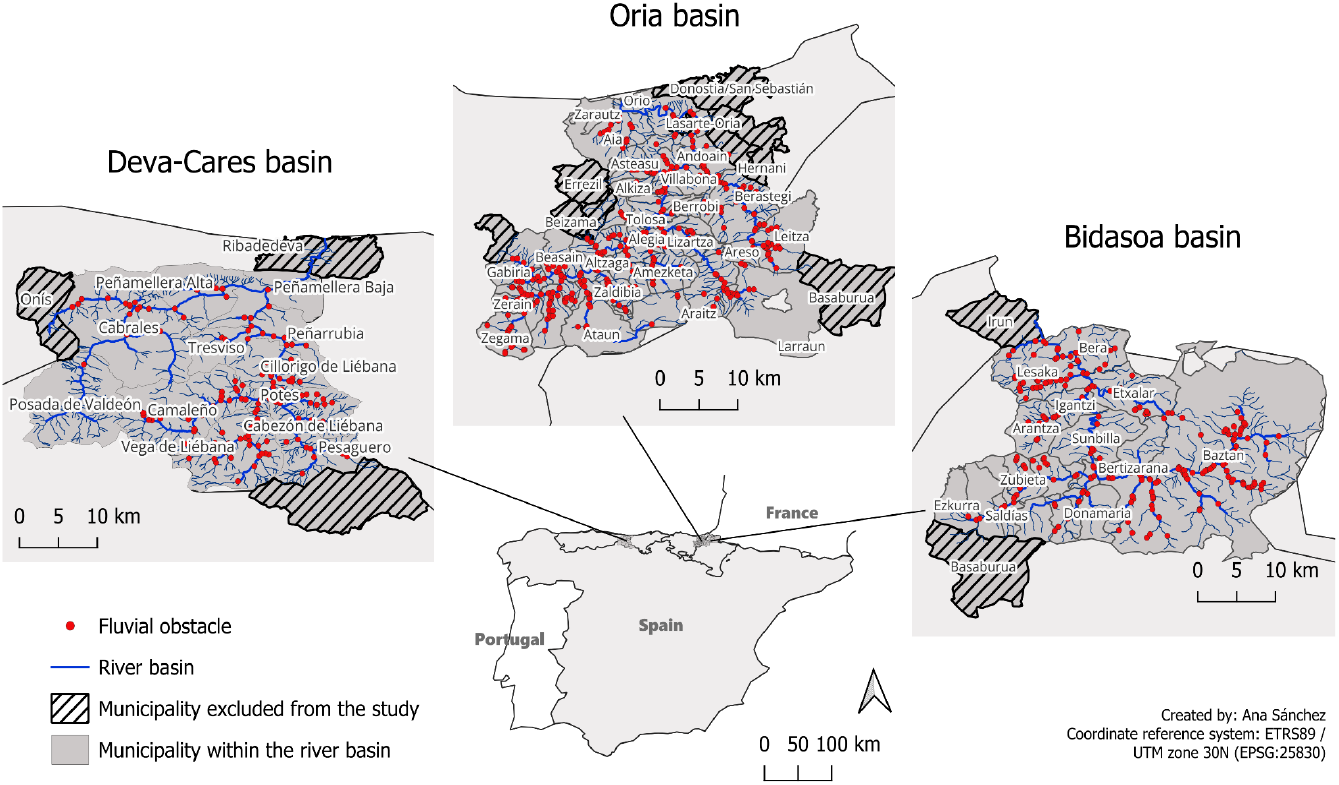
Location of the three river basins in which the telephone surveys were conducted. The target basins are located in four communities: Asturias, Cantabria, the Basque Country, and Navarra.

These basins are among the very few areas of the Iberian Peninsula that still hold natural populations of Atlantic salmon (*Salmo salar* Linnaeus, 1758), a species of considerable ecological and fishery relevance but currently in persistent decline, with habitat restoration being one of the most effective but controversial measures for its recovery (Horreo et al., 2011; Clavero et al., 2025; García-Vega et al., 2025).

This area presents a high density of fluvial obstacles, according to regional databases and the European AMBER Barrier Atlas (AMBER Consortium, 2020; Izeta-Zalduendo et al., in prep).

### Sampling design and collection

The target population for the survey comprised adults residing in municipalities within these catchments that have at least one fluvial obstacle in the river network that flows through them. According to this last criterion, some headwater and river mouth towns were excluded (Figure 1).

An intended sample of 1,200 respondents was established, 400 for each basin, a figure statistically significant at the 95% confidence level according to the most recent population census for the area (National Institute of Statistics (INE, 2024). The sample also held equal representation of each gender and similar proportions across the four established age groups (18–34, 35–49, 50–64, >65).

From the total number of respondents, the present study focuses on the 180 individuals who self-reported being anglers. To answer the question about whether anglers’ perceptions differ from those of the public, we compared that group with a subsample of 180 people who do not practise this activity. To reduce the possible influence of other sociodemographic factors, we applied a stratified sampling that kept the same gender, age, and political ideology ratios in both groups, as previous studies have described correlations between these aspects and people’s opinions on environmental issues (Diessner et al., 2020; Dopico et al., 2022).

### Questionnaire design

The study employs an *ad hoc* questionnaire tailored to the specific context and goals of the study. The questionnaire design was based on insights from academic literature and media sources, plus the contributions received from eight semi-structured interviews with diverse stakeholders from the study area, including members of the administration’s environmental department, environmental groups, residents of riverside areas, and anglers. A panel of 16 experts from diverse areas (zoology, ecology, data management, hydrology, sociology, psychology, activism, fishing, and local entities) validated the clarity and content of the initial draft. Since a pilot study was unfeasible due to territorial limitations, the internal consistency of the questionnaire items was tested using the responses provided by the final respondents and deemed acceptable (Cronbach’s alpha = 0.73).

The final questionnaire consisted of three sections: (i) a first section collecting general opinions about fluvial obstacles and their impacts; (ii) a second section addressing specific dams and weirs in the vicinity of the respondent; and (iii) a final section gathering sociodemographic parameters, including gender, age, socioeconomic status, hobbies, political and religious ideology, and level of connection with nature. The latter parameter was assessed using a translated version of two selected items from the scale proposed by Gagnon Thompson & Barton (1994), chosen for their representativeness, as the full scale was considered too lengthy and no other viable alternatives suitable for telephone administration were identified.

Following recommendations provided by Dopico et al. (2022), an effort was made to avoid overlooking small structures - such as weirs, ramps, culverts, and fords - which actually constitute 85% of barriers in European rivers but are too often underrepresented in studies on river fragmentation. This inclusion was made explicit in a short text that interviewers read to respondents before starting, emphasizing that the survey items refer not only to dams but also to other smaller obstacles.

The questionnaire was designed for telephone administration. Since the survey gathered no personal data, participants’ verbal consent was deemed enough to enable data collection and processing. The study received approval from the University of Navarra’s Research Ethics Committee (reference number 2024.286) before starting the survey.

This paper focuses exclusively on the analysis of the answers given to the initial part of the questionnaire (12 items available in Supplementary material), taking into account the socio-demographic parameters collected in the last part. The items in this first section assess social perception of barriers’ environmental impacts, and they also try to evaluate the social acceptance of the most prominent discourses and myths surrounding resistance to obstacle removal (Jørgensen & Malm, 2013; Fox et al., 2016; Hommes, 2022). As recommended in previous research (Diessner et al. 2020), the survey not only included items to gather opinions but also questions designed to assess actual knowledge about dam-related issues, which may help guide public communication efforts on removal processes.

## Results

### Sample characterization

The angler sample is predominantly male (76.7% compared with 23.3% female), spans a wide age range, and is evenly distributed across the three study areas. It shows low to medium educational attainment and a medium income level. Its political ideology tends towards the left, and most respondents adhere to the Catholic religion (Table 1).

**Table 1.** Sociodemographic characterisation of the respondent samples used in the study. Most frequent replies are bold.

|  |  | Anglers |  | Non-anglers |  | p-value |
| --- | --- | --- | --- | --- | --- | --- |
|  |  | n | % | n | % |  |
| Gender | Male | **138** | 76.7 | **136** | 75.6 | 0.806 |
|  | Female | 42 | 23.3 | 44 | 24.4 |  |
| Age | 18-30 | 33 | 18.3 | 33 | 18.3 | 0.999 |
|  | 31-44 | 48 | 26.7 | 47 | 26.1 |  |
|  | 45-64 | **57** | 31.7 | **58** | 32.2 |  |
|  | 65+ | 42 | 23.3 | 42 | 23.3 |  |
| Residential area | Deva-Cares basin | 56 | 31.1 | 70 | 38.9 | 0.197 |
|  | Oria basin | 59 | 32.8 | 59 | 32.8 |  |
|  | Bidasoa basin | 65 | 36.1 | 51 | 28.3 |  |
| Level of education | Very low | 4 | 2.2 | 10 | 5.6 | 0.011* |
|  | Basic | **60** | 33.3 | 34 | 18.9 |  |
|  | Medium | 49 | 27.2 | 44 | 24.4 |  |
|  | Technical | 38 | 21.1 | **49** | 27.2 |  |
|  | Superior | 29 | 16.1 | 42 | 23.3 |  |
|  | DK/NA | 0 | 0 | 1 | 0.6 |  |
| Employment situation | Housekeeper | 1 | 0.6 | 7 | 3.9 | 0.181 |
|  | Unemployed | 12 | 6.7 | 17 | 9.4 |  |
|  | Retired | 47 | 26.1 | 49 | 27.2 |  |
|  | Student | 8 | 4.4 | 6 | 3.3 |  |
|  | Currently working | **112** | 62.2 | **101** | 56.1 |  |
| Income level | Low | 6 | 3.3 | 8 | 4.4 | 0.05 |
|  | Medium | **120** | 66.7 | 73 | 40.6 |  |
|  | High | 13 | 7.2 | 18 | 10 |  |
|  | DK/NA | 41 | 22.8 | **81** | 45 |  |
| Political ideology | Extreme left | 46 | 25.6 | 44 | 24.4 | 0.989 |
|  | Left | **49** | 27.2 | **50** | 27.8 |  |
|  | Center | 35 | 19.4 | 35 | 19.4 |  |
|  | Right | 21 | 11.7 | 24 | 13.3 |  |
|  | Extreme right | 17 | 9.4 | 14 | 7.8 |  |
|  | DK/NA | 12 | 6.7 | 13 | 7.2 |  |
| Religious affiliation | Practising Catholic | 66 | 36.7 | 48 | 26.7 |  |
|  | Non-practising Catholic | **79** | 43.9 | **83** | 46.1 |  |
|  | Believer of another religion | 6 | 3.3 | 16 | 8.9 |  |
|  | Agnostic | 3 | 1.7 | 9 | 5 | 0.066 |
|  | Indifferent, non-believer | 23 | 12.8 | 17 | 9.4 |  |
|  | Atheist | 2 | 1.1 | 5 | 2.8 |  |
|  | DK/NA | 1 | 0.6 | 2 | 1.1 |  |
| Level of connection with nature<br>A: The most important reason for nature conservation is human survival | Strongly disagree | 5 | 2.8 | 9 | 5.0 |  |
|  | Disagree | 2 | 1.1 | 16 | 8.9 |  |
|  | Neutral | 25 | 13.9 | 30 | 16.7 | 0.020* |
|  | Agree | 17 | 9.4 | 26 | 14.4 |  |
|  | Strongly agree | **78** | 43.3 | 43 | 23.9 |  |
|  | DK/NA | 53 | 29.5 | **56** | 31.1 |  |
| Level of connection with nature<br>B: Human beings are part of the ecosystem, just like other animals | Strongly disagree | 5 | 2.8 | 5 | 2.8 |  |
|  | Disagree | 0 | 0.0 | 5 | 2.8 |  |
|  | Neutral | 24 | 13.3 | 27 | 15.0 | 0.249 |
|  | Agree | 23 | 12.8 | 33 | 18.2 |  |
|  | Strongly agree | **76** | 42.2 | **55** | 30.6 |  |
|  | DK/NA | 52 | 28.9 | 55 | 30.6 |  |

DK/NA: don't know / no answer
\*Differences were considered statistically significant at p-values below 0.05

Gender, age, and political ideology were taken as key opinion-predictor factors when creating a comparable subsample of non-anglers, ensuring that the differences observed between the two groups could not be attributed to variations in these factors. The remaining sociodemographic variables of the two samples were compared using chi-square tests or one-way ANOVA, depending on the type of variable, without taking into account the DK/NA answers. No statistically significant differences were detected, with the sole exception of educational attainment, and one of the items related to the assessment of connection with nature.

### Questionnaire results

In the following section, the term *dam* is used broadly to also encompass other types of fluvial barriers of a different nature.

#### Reported knowledge

Although the majority of interviewees reported having “little” to “medium” knowledge about river dams (median=5, on a scale from 1 to 10), the number of anglers reporting high or very high levels of knowledge is significantly greater than that of non-anglers (Figure 2.A).

**Figure 2.**
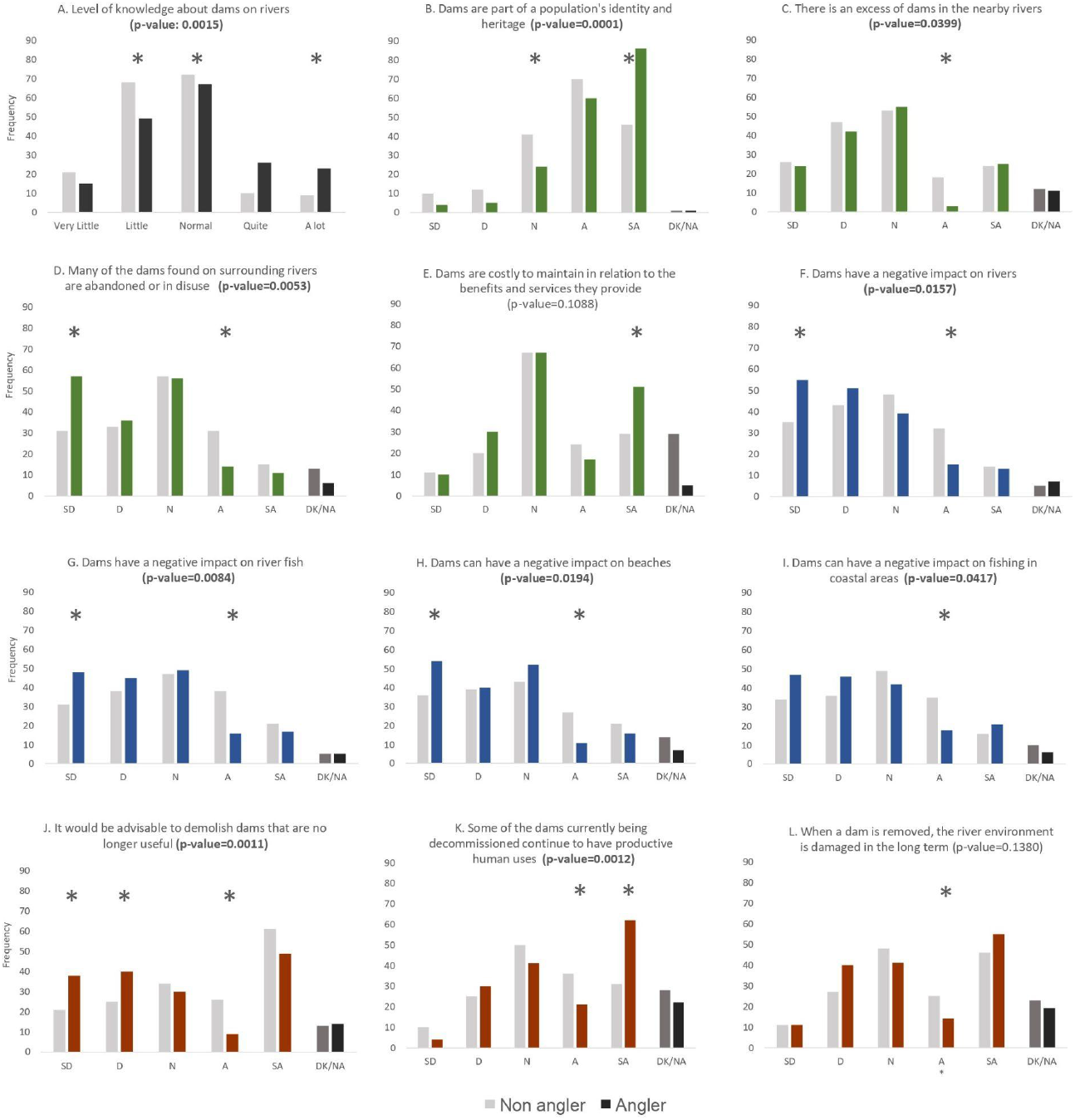
Responses provided by respondents to each statement (n= 360). Answers to item (A) were registered on a scale from 1 to 10. The figure shows results grouped in five categories, as follows: 1-2=very little; 3-4=little; 5-6=normal; 7-8=high; 9-10=very high SD: strongly disagree, D: disagree, N: neutral, A: agree, SA: strongly agree, DK/NA: don’t know /no answer * = Significant difference between groups (p < 0.05) Different color palettes show thematic grouping of items: Reported knowledge (A); Dams in the landscape (B, C, D, E); Dam impacts (F, G, H, I); Dam removal (J, K, L)

#### Dams in the landscape

Both groups tend to see dams as part of a population’s identity and heritage, but anglers’ agreement with this statement is stronger than non-anglers’ (Figure 2.B). Something similar can be observed when participants were asked about the number and abundance of dams (Figure 2.C, D): although both groups seem to have a similar response pattern, in this case, non-anglers report a greater belief that there is an excess of dams in their surrounding rivers and that many of them are abandoned or in disuse.

The two groups show a noticeable proportion of respondents who adopt a neutral position regarding the cost–benefit ratio of dams (Figure 2.E), although the number of anglers who think that dams are costly to maintain in relation to their benefits is significantly higher than that of non-anglers. Also, the higher number of DK/NA responses among non-anglers is noticeable, a case that can be observed in most of the questions.

#### Dam impacts

Both groups tend to be either neutral or to disagree with the statement that dams cause negative impacts on rivers, river fish, beaches or coastal fishing, although anglers’ disagreement is significantly stronger than that of non-anglers (Figure 2.F, G, H, I).

#### Dam removal

Opinions on the advisability of demolishing unused dams (Figure 2.J) show a bimodal distribution, with a large percentage of respondents supporting demolishing dams that are no longer useful and a smaller but noticeable group opposed. The proportion of anglers who disagree with that statement is significantly larger than that of non-anglers (47% of anglers disagree or strongly disagree versus 27.5% of non-anglers).

Regarding one of the most widespread pieces of misinformation on the subject (Figure 2.K), whereas non-anglers’ responses show an almost Gaussian distribution, in the case of anglers, there is a significant proportion of respondents who strongly agree with the statement that many of the dams currently being decommissioned still have productive human uses.

The responses to the perception of dam removal causing long-term negative effects (Figure 2.L) do not show significant differences between groups, which follow a bimodal distribution. In both of them, there is a great percentage of respondents who deem dam removal to cause negative impacts on the environment.

## Discussion

The gender composition of the anglers group in our study (76.7% males vs. 23.3% females) differs notably from that of the survey sample (50% females and 50% males) and that of the area’s population (50.5% females and 49.5% males) (INE, 2024). This result aligns with previous studies reporting greater participation in recreational fishing among males than females, both in Spain and in other countries (Morales-Nin et al. 2025). Actually, the number of males in our study was more than three times greater than the number of females, a greater disproportion than that reported by Morales-Nin et al. (2025).

Regarding the level of connection with nature, the results obtained to assess respondents’ anthropocentric or ecocentric perspectives did not align with expectations. The two items of the questionnaire were taken from a more comprehensive scale by Gagnon Thompson & Barton (1994). According to the authors, individuals who disagree with statement A (“The most important reason for nature conservation is human survival”) are expected to agree with statement B (“Human beings are part of the ecosystem just like other animals”), and vice versa. However, a general tendency was observed in both groups to agree with both affirmations (Table 1. This pattern suggests that the selection of items, or their use out of the whole scale, may not have been fully appropriate for identifying ecocentric/anthropocentric positions in the present context. Actually, when used in an isolated way (as we did) and not as part of the scale, an agreement with both statements may not necessarily be contradictory: a person may see the human species as part of the ecosystem and also prioritize our needs in any action we take, just as any other species does. In any case, considering the uncertainty of how to correctly interpret these answers, although these items may still provide some relevant information, they were given limited weight in characterizing the samples and in interpreting the study results.

The survey responses show some general patterns of interest. On the one hand, the number of DK/NA answers is almost always higher in the control group than among anglers. Besides, the proportion of respondents who manifest strong positions (strongly agree, strongly disagree) is usually higher in anglers than in non-anglers. These two observations point out that, in this study, anglers show a greater tendency to express clear and sometimes extreme positions regarding dams than non-anglers. On the other hand, we observed a high proportion of neutral responses in both groups across several items. The actual meaning of the neutral position in a Likert scale has been disputed in the literature, as respondents may not interpret and use such midpoint in the way that survey developers intended (Chyung et al. 2017; Garland, 1991; Nadler et al. 2015). Some authors have raised concerns that respondents may use such a midpoint as a “dumping ground” when answering unfamiliar items, especially if they perceive that option as more socially desirable than showing a lack of opinion (Chyung et al. 2017). In our study, both groups reported low to moderate levels of self-perceived knowledge about river barriers. This lack of specific knowledge may therefore be related to the lack of positioning that such a high percentage of neutral responses show.

Despite that general result, when comparing both groups, we see that anglers report higher knowledge than non-anglers. However, their positioning with regard to other statements does not suggest that they are more aware of the negative environmental impacts caused by river barriers than the other group, not even of those impacts that might be more readily perceptible through their direct experience, such as effects on the river or on fish populations. In fact, according to our data, anglers’ disagreement with the statement that dams cause negative ecological effects is significantly stronger than that of non-anglers (Figure 2.F).

A factor that could be influencing these responses is the lower level of educational attainment reported by anglers compared with the control sample -a result consistent with literature associating formal education with a reduced likelihood of practising freshwater fishing (Arlinghaus et al. 2021)-. In this line, a study recently carried out by Banha and collaborators (2024) associated higher awareness of non-native freshwater fish species to higher levels of education. However, the broader literature does not consistently identify this sociodemographic aspect as a key determinant in shaping perceptions, since there are studies that relate higher education levels with greater awareness about water issues (Gilg & Barr, 2006; Meyer, 2015; Sanchez et al. 2016), whereas others do not find such an effect (Diessner et al., 2019; Diessner et al., 2020; Taylor and Signal 2016). According to literature, there are several other personal and social factors - such as childhood experience, personality, or place attachment - that may influence environmental concern (Gifford & Nilsson, 2014) and were not registered in this survey.

Besides sociodemographic differences, other factors may influence anglers’ perception of river barrier impacts in the ecosystem. Anglers’ angling satisfaction is highly catch-dependent (Arlinghaus 2006; Gundelund et al. 2022). More specifically, catch rate and size of captures are two main drivers of satisfaction with catch in recreational fishers (Beardmore et al. 2015; Birdsong et al. 2021). This effect may be especially relevant in trophy-oriented anglers, who represent around 20% of recreational fishers in some European areas (Bonnichsen et al. 2019). Some species, such as wels catfish *Silurus glaris*, have more trophy potential because they grow larger than others, and may therefore be preferred by trophy-oriented anglers (Arterburn et al. 2002). In European countries, the wels catfish is quite popular among recreational fishers, also in places where it is non-native (Fromherz et al. 2024). Actually, a recent study in Portugal reported that catfish anglers perceived the species as positive not just for angling, but also for other fish (Gago et al. 2025). The presence of river barriers favours wels catfish, as it grows bigger in large rivers (not present in our study basins) and in dammed waters. Impounded water may also favour the trapping and proliferation of other species of particular interest or appeal to some groups of anglers (Kruk & Penczak, 2003; Alexandre & Almeida, 2010; Clavero et al. 2011; Clavero et al. 2013). It may derive from this that trophy-oriented anglers in our region perceive river barriers as beneficial to the fish they seek, and therefore not so damaging to the environment. This same reason may be behind the result of more anglers than non-anglers disagreeing with the idea of many nearby dams being abandoned or in disuse (Figure 2.D): those anglers who look for fish in reservoirs actually use dams for that end, whereas people with no interest in that activity may not even think of this as a service these structures provide.

Besides catch, diverse fisher profiles share other motives such as escapism (being outdoors), challenge (Magee et al. 2018), and the ambiguity of the fishing result (i.e. the uncertainty of the catch; Arlinghaus 2024). These motives can be easily fulfilled and therefore drive satisfaction in anglers. If a certain environment meets the expectations of a visitor, that person may be less prone to appreciate the downsides of that feature s/he’s enjoying, or to see them as relevant.

Not all anglers are interested in fish that grow in impounded waters. Some native migrant species in the area (such as Atlantic salmon *Salmo salar*, and brown trout *Salmo trutta*) prefer current waters and are negatively affected by river barriers. The existence of two clearly different types of angling - one that benefits from the existence of dams, and another kind that needs stretches of free-flowing rivers - may help explain the bimodal trends observed in some of the results obtained. For instance, respondents who expressed disagreement with the removal of abandoned dams (Figure 2.J) may do so because their fishing activity benefits from the existence of such structures. In contrast, those who support their demolition may be associated with other fishing modalities, related to native species that are strongly affected by river fragmentation. This information was not collected in the present study, but our results point out that it would be interesting to explore this aspect in future studies with similar populations.

Direct contact with nature is thought to raise environmental awareness (Rosa and Collado, 2019), and therefore we would expect anglers to show greater knowledge or sensitivity towards ecological impacts caused by dams. However, our results show that they tend to disagree with the idea of river barriers causing negative impacts on fish and water ecosystems. This may be partially explained by the reasoning stated above regarding trophy-oriented anglers. Also, the outcomes of contact with nature are not the same for all people, and the angling community shows a diversity of opinions regarding river ecosystems, even if all its members spend a considerable amount of time in direct contact with this ecosystem (Eden and Bear, 2011). However, the scientific literature shows that recreational fishing can foster environmental stewardship (Shephard et al. 2024), and that recreational fishers identify environmental values that align with those of conservationists (Young et al. 2016). In our study, we find an example of the latter in the high percentage of anglers who agree with the ecocentric statement that “humans are part of the ecosystem just like other animals” (Table 1). This lays common ground for building consensus between anglers and conservationist stakeholders, and provides useful information to design effective communication and education campaigns.

According to our results, anglers seem more susceptible than the control group to believing some widespread misconceptions about river barrier removal. In both groups there is a noticeable proportion of respondents who believe that part of the barriers currently being removed are still serving productive uses (Figure 2.K), even if, under Spanish law, only those whose concessions have expired may be dismantled (Royal Decree 849/1986, Royal Legislative Decree 1/2001). This result may be partially explained by the fact that in many cases recreational fishing takes place in or benefits from dams, weirs and other barriers that have no other uses. Participants of this activity may therefore perceive these structures as still functional, whereas non-anglers deem them abandoned or not in use. Also, illegal uses ーusually known, even if not approved, by regular visitors of a placeー may still be happening in abandoned barriers, and thus make them look as active even if official registers say the contrary.

A high number of respondents also think that dam removal causes long-term damage in river environments (Figure 2.L), even though several studies demonstrate that rivers are highly dynamic and resilient ecosystems that rapidly recover from damage (Bednarek 2001; Bellmore et al. 2019). It is noticeable that in both samples there is a higher proportion of DK/NA answers than in other items (Figure 2.K and L), which may denote that respondents are not as familiar with these narratives as the literature suggests (Lejon et al., 2009; Hommes, 2022). These results indicate that, although a non-negligible proportion of the population appears to be aware of and informed about the impacts caused by dams and other barriers, the topic remains largely unfamiliar to the public included in this study. Awareness-raising campaigns therefore continue to be necessary to counter misinformation and to publicise both the impacts generated by fluvial barriers and the benefits of their removal, in order to achieve successful removal processes supported by citizens. In line with this, different dam acceptance emphasizes the need to involve local citizens and stakeholders in decision-making processes related to water management.

The views of anglers ーwho are closely connected to fluvial ecosystemsー cannot be overlooked, as river restorations, such as dam removal interventions, will rarely achieve social sustainability without local support. Moreover, recreational fishers hold a stewardship potential that can maximize biodiversity conservation efforts (Shephard et al. 2023), and their ecological knowledge is increasingly getting recognized and valued in academic circles (Granek et al. 2008; Löki et al. 2023). Getting to know their perception on water-related issues, such as river barrier removal, is paramount to tackle such processes in a way that minimizes conflicts and opens room for understanding and cooperation. Taking this information into account when designing participatory or information processes will help build engagement among stakeholders, and increase projects’ chances of success. In a time when restoring stream connectivity is gaining momentum, combining social criteria with the ecological and economic focus seems more urgent than ever.

## Supporting information

Supplementary - Questionnaire English

## Acknowledgements

This study is part of a broader project entitled “ConnectFish: analysis of the ecological connectivity of dams in relation to the conservation status of Iberian fish species through a multidisciplinary approach”. This work was funded by the Biodiversity Foundation of the Ministry for the Ecological Transition and the Demographic Challenge of Spain (MITECO), within the framework of the Recovery, Transformation and Resilience Plan (PRTR), financed by the European Union – NextGenerationEU. The opinions and conclusions expressed in this publication are the sole responsibility of the persons who author them, and do not necessarily reflect the views of the entities that financially support the project. Francisco J. Peñas is supported by a Ramón y Cajal Grant (ref. RYC2024-049751-I) from the Ministry of Science, Innovation and Universities. Alexia M. González is supported by a Ramón y Cajal Grant (ref. RYC2023-045780-I) from the Ministry of Science, Innovation and Universities. The researchers would like to thank the stakeholders who contributed to the development of the questionnaire by giving their time to participate in several interviews, as well as the panel of experts who subsequently reviewed and validated it.

## Data availability statement

A version of the survey used, translated into English, can be found in Supplementary material.

The dataset containing the survey responses associated with this manuscript is available at the following DOI: https://doi.org/10.6084/m9.figshare.30800141

All the data about the research project to which this study belongs can be accessed at the following website: https://bit.ly/ConnectFish

