## Supplementary - Questionnaire English for "Do anglers take the bait? Anglers’ perceptions about fluvial barriers in three river basins in Northern Spain"

### UNIVERSITY OF NAVARRA

#### Analysis of Social Perception of Dams, Their Impacts and the Ecosystem Services They Provide.

This survey is part of the project entitled "Analysis of the Ecological Connectivity of Dams in Relation to the Conservation Status of Iberian Fish: A Multidisciplinary Approach". Its aim is to study Spanish society's views on river dams (including other artificial water barriers such as weirs, dykes, and impoundments), their impacts, and the ecosystem services they provide. The study is funded by the Biodiversity Foundation and carried out by a research team from the University of Navarra in collaboration with the Environmental Hydraulics Institute Foundation of Cantabria. The lead researchers are Ana Villarroya and Ana Sánchez, who may be contacted at if needed. Participation in this survey is entirely voluntary, and it is estimated that it will take approximately 7 minutes to complete. By completing this questionnaire, you expressly give your consent to the University of Navarra for the use of the information. The data collected are anonymous and will be used exclusively for research purposes.

To start, I would like to ask whether you are:

a) Man
b) Woman
c) Other

Could you indicate your year of birth? ________

In which municipality do you reside? Dropdown with the localities included in the project

GENERAL QUESTIONS

1) On a scale of 1 to 10, where 1 is "very little" and 10 is "a lot", how would you rate your level of knowledge about river dams?

9) Please indicate your level of agreement with the following statements on a scale from 1 (strongly disagree) to 5 (strongly agree). (Rotate items)

a) Dams are part of the identity and heritage of a community.
b) There are too many dams in the nearby rivers.
c) Many dams in your area are abandoned or out of use.
d) Dams are costly to maintain relative to the benefits and services they provide.

e) Dams have a negative impact on rivers.
f) Dams have a negative impact on river fish.
g) Dams can negatively affect beaches.
h) Dams can negatively affect coastal fishing.
i) It would be advisable to remove dams that no longer have any use.
j) Some dams currently being removed still have productive human uses.
k) When a dam is removed, the river environment is left damaged in the long term.

SOCIO-DEMOGRAPHIC QUESTIONS

33) Indicate your level of agreement with the following statements (1 = strongly disagree, 5 = strongly agree):

a) The most important reason for nature conservation is human survival.
b) Human beings are part of the ecosystem just like other animals.

34) What is the highest official level of education you have reached, regardless of whether you finished it or not?

a) No formal education
b) Incomplete primary education (schooling up to approximately age 10)
c) Completed primary education (Basic General Education, compulsory schooling up to ages 14–16)
d) Secondary education (Baccalaureate or equivalent, up to approximately age 18)
e) Vocational training (Intermediate or advanced vocational qualifications)
f) University studies – first cycle (e.g., Diploma or equivalent)
g) University studies – second cycle (e.g., Bachelor’s degree)
h) University studies – third cycle (e.g., Master’s degree, PhD)

35) What is your current occupation?

a) Currently employed
b) Unemployed (previously employed)
c) Seeking first job
d) Retired, pensioner, or permanently disabled
e) Student (not working)
f) Homemaker

36) Regarding your leisure preferences… Which of the following outdoor activities do you practice regularly?

a) Hiking
b) Hunting
c) Angling
d) Water sports (rowing, canoeing…)
e) Photography
f) Other: _______

37) What is the total net monthly income of all members of your household?

a) Up to €1,000 per month
b) €1,001–€1,999 per month
c) €2,000–€2,999 per month
d) €3,000–€3,999 per month
e) €4,000–€4,999 per month
f) €5,000–€5,999 per month
g) €6,000 or more per month
h) Prefer not to say

38) When talking about politics, people often refer to "left" and "right". Using a scale from 1 to 10, where 1 means "the most left-wing" and 10 "the most right-wing", where would you place yourself?

39) How would you define yourself in terms of religion?

a) Practicing Catholic
b) Non-practicing Catholic
c) Believer of another religion
d) Agnostic (do not deny the existence of God, but do not affirm it either)
e) Indifferent, non-believer
f) Atheist (deny the existence of God)
g) Prefer not to answer

THANK YOU VERY MUCH FOR YOUR PARTICIPATION
